# A high-throughput multiplex array for antigen-specific serology with automated analysis

**DOI:** 10.1101/2023.03.29.534777

**Authors:** A.F Rosenberg, J.T. Killian, T.J. Green, J. Akther, M.E. Hossain, Q. Shang, S. Qiu, Y. Guang, T.D. Randall, F.E. Lund, R.G. King

## Abstract

**High-throughput customizable CBA automated script-based analysis:** The utility of high-throughput systems to evaluate antigen-specific antibody (Ab) has been highlighted by the SARS-CoV-2 pandemic. Pathogen-specific Ab levels are often used to assess protection following vaccination and, in the case of novel pathogens, an indication of prior exposure. Several platforms exist to visualize antigen-specific Ab, however most are not quantitative and are difficult to scale for population level studies. Additionally, the sensitivity across platforms differs making direct comparisons between studies difficult. Cytometric bead arrays (CBA) are an attractive platform for antigen-specific Ab measurements as they can be used to assess Ab reactivity against several antigens and of several isotypes to be performed simultaneously. Additionally, CBAs exhibit high sensitivity and can be designed to provide quantitative measurements. Using commercially available particles, a biotin-Streptavidin bead loading strategy, and the inclusion of indirect standards, we describe a flexible system that can be modified to include a variety of antigens. Here we describe two arrays, focused on antigens derived from either β-coronaviruses or influenza virus. To support the high-throughput capacity of this system, we developed a suit of automated software tools, the CBA Toolbox, to process raw data into antigen-reactive IgM, IgA, and IgG concentrations. We describe quality control requirements, assay performance, and normalizations to accurately quantitate antigen-specific Ig.

## 1. Introduction

Pathogen-specific serum antibody (Ab) levels are commonly used to determine prior exposure history, to measure vaccine responsiveness, and to predict whether someone who was previously infected or vaccinated will be protected from reinfection. The COVID pandemic has highlighted the need for rapid and accurate quantification of antigen-specific Abs in human biologic samples. Early in 2020, the human immune response to SARS-CoV-2 was uncharacterized and considerable effort was devoted to determining how individuals respond to infection and whether particular Ab responses correlated with disease outcomes. Even following the approval of multiple SARS-CoV-2 vaccines, it was still unclear which serological features of the humoral immune response afforded protection against infection or re-infection with SARS-CoV-2. Unfortunately, the evolution of the Sars-2-CoV-2 pandemic over the last three years has illuminated the shifting landscape of viral variants and the uneven protection afforded by the original vaccination formulations [1, 2]. Despite numerous published large cohort studies [3-6] that addressed the magnitude and longevity of SARS-CoV-2 humoral immunity, it is clear that there is still a need to develop inexpensive, high-throughput methodologies to evaluate Ab responses following infection or vaccination [7, 8]. Additionally, it is now increasingly obvious that not all Abs directed toward the SARS-CoV-2 spike protein (C SP) are protective and that we need quantitative Ab detection platforms that can measure multiple isotypes of C SP Abs and can assess C SP Ab specificity for Spike variants and sub-domains of C SP. Although several platforms to detect SARS-CoV-2-reactive Abs [9–16] have been described, few of these platforms provide quantitative data. Thus, cohort data, which was generated using the different Ab platforms, cannot be directly compared across studies. This limitation has hampered our ability to define specific concentrations of C SP reactive Abs that correlate with protection. Such quantitative analyses are becoming increasingly important as SARS-CoV-2 variants that are resistant to prior vaccine responses continue to emerge [2]. Additionally, there are very limited open-source, non-instrument-linked options for automated analysis that would be suitable for high-throughput, batched Ab reactivity profiling [17]. Although the COVID pandemic has increased our appreciation for the utility of platforms that can be used to evaluate antigen-specific Ab, such platforms are important for many other pathogens including seasonal influenza (INF) virus. To address this need, we developed a highly customizable cytometric bead array (CBA) platform to quantitate specific Abs of IgG, IgM, and IgA subclasses against several pathogen associated antigens. Coupled with an automated MATLAB analysis pipeline, the CBA Toolbox that is highly configurable like the assay itself, this system allows for the high-throughput quantification of Abs reactive to a wide variety of antigen panels.

CBAs have been widely employed to determine the concentration of diverse analytes from biological samples, including Abs [18]. CBAs offer several advantages over traditional ELISA-based approaches including enhanced sensitivity and a broad dynamic range. In addition, CBA ligands can be multiplexed, thereby allowing for analysis of many antigens simultaneously in a high-throughput format [14, 18].

Here we describe development of a highly customizable CBA platform that, when coupled with automated MATLAB analysis pipeline, can be used in a high-throughput manner to rapidly quantitate IgG, IgM, and IgA Abs specific for a variety of pathogen-derived antigens. One CBA consisted of 12 antigens derived from β-Coronavirus (β-CoV) virus spike ectodomains (SP) and nucleocapsid proteins (NP). The second array incorporated 14 INF hemagglutinins (HA) in addition to influenza NP and non-structural protein 1 (NS1). To support this detection platform, we developed a post-acquisition, MATLAB-based analytic pipeline that allowed for streamlined data analysis and enabled high-throughput use. In addition, we evaluated the performance of the array using panels of monoclonal and polyclonal human Abs. Using these reagents, we determined antigen-specific correction factors allowing for the accurate quantification of serum Ab of the IgM, IgA, and IgG subclasses toward each of the array antigens. Although the configuration of the arrays described here was focused on β-CoV- and INF-derived antigens, the platform is flexible and utilizes indirect standards, which allow for quantitation of Ab reactivity to essentially any protein antigen.

## 2. Material and methods

### 2.1 Array generation

Streptavidin (SA) (SouthernBiotech 7105-01) was buffered exchanged into PBS using PD-10 columns (Cytiva 17085101) and then diluted to 2mg/mL. SA was then conjugated to spherotech 4μm and 5μm Carboxy Blue Particle Array Kits (Spherotech CPAK-4067-8K and CPAK-5067-10K) using 1-Ethyl-3-(3-dimethylaminopropyl)carbodiimide (EDC) chemistry. For conjugation, 1e8 spherotech particles were isolated by centrifugation at 10,000Xg for 3 min. After carefully removing the supernatant, the bead pellet was resuspended in 0.5 mL SA in PBS. Following complete resuspension by pipetting, 0.5 mL of 6mM EDC, dissolved in 0.05M MES buffer pH 5.0, was added, and the reaction mixture was rotated at room temperature (RT) overnight (O/N). After the conjugation reaction was complete, 0.1 mL 1M tris pH 8.0 was added to quench the reaction. Following a 1 hr incubation, the beads were harvested by centrifugation as described above and washed twice in 1mL PBS. Following the final wash, beads were resuspended at 10e8 particles/mL in PBS with 0.25% NaN3 and stored at 4°C.

Following SA conjugation, quality control experiments were performed to determine the degree and uniformity (when multiple particle sizes and/or peak identities are used) of labeling by staining the SA-conjugated particles with Alexa488-labeled-biotinylated hemagglutinin (HA) from the H1 HA derived from influenza A/PR/8/34 (PR8). Individual array constituents were mixed and diluted to 1e6 of each particle/mL. 40 µL of serial dilutions of PR8 HA were prepared in a 96 well U bottom plate (Corning 3797) ranging from 1µg/mL to 2ng/mL. 5µL of the bead suspension was added, mixed by pipetting, and incubated for 15 min at RT. 200µL PBS was added and the plate was centrifuged at 3000Xg for 5min. The beads were resuspended in 80µL PBS. PR8 HA bound to SA conjugated beads was visualized by flow cytometry.

Following SA coupling and quality control analyses described above, biotinylated recombinant antigens were passively absorbed onto the individual particles. For antigens described here (Table 1), a single biotin was added enzymatically onto a carboxy terminal AVI tag. SA conjugated particles were harvested by centrifugation as described above and resuspended in 1mg/mL of the recombinant proteins in 1% BSA in PBS. Antigen loading was carried out by rotating overnight at 4°C. Following absorption, the beads were harvested by centrifugation as described, and washed twice with 1% BSA in PBS. Finally, the antigen coated beads were resuspended at 1e8 particles/mL 1% BSA in PBS, 0.25% NaN3 and stored at 4°C until used.

**Table 1.** List of antigens used to generate the Inf and β-CoV antigen Arrays. The size and peak number of each array element is listed, with the biotinylated recombinant protein loaded to generate the two arrays. The protein accession numbers are provided.

|  |  |  |  |
| --- | --- | --- | --- |
| **β-CoV** |  |  |  |
| **4um-2** | S NP | Sars-CoV-1 Nucleocapsid Protein | AAP13445.1 |
| 4um-3 | M NP | MERS Nucleocapsid Protein | P_009047211.1 |
| 4um-4 | O NP | OC43 Nucleocapsid Protein | KR055606.1 |
| 4um-5 | H NP | HKU1 Nucleocapsid Protein | Q5MQC6.1 |
| 4um-6 | C NP | Sars-CoV-2 Nucleocapsid Protein | QHD43423 |
| 5uM-1 | S SP | Sars-CoV-1 Spike ectodomain | AAX16192.1 |
| 5uM-2 | S RBD | Sars-CoV-1 Receptor binding domain | AAX16192.1 |
| 5uM-3 | C SP | Sars-CoV-2 Spike ectodomain | QHD43416.1 |
| 5uM-4 | C RBD | Sars-CoV-2 Receptor binding domain | QHD43416.1 |
| 5uM-5 | C NTD | Sars-CoV-2 N-Terminal domain | QHD43416.1 |
| 5uM-6 | M SP | MERS Spike ectodomain | AKI29255.1 |
| 5uM-7 | O SP | OC43 Spike ectodomain | AAT84362.1 |
| 5uM-8 | H SP | HKU1 Spike ectodomain | Q5MQD0.1 |
| 5uM-9 | CA09 | Influenza HA (H1) | NC_026433.1 |
| **Influenza** |  |  |  |
| **Peak #** | **Antigen** | **Antigen description** | **Protein Accession #** |
| 5uM-1 | PR8 | Influenza HA (H1) | MH785011.1 |
| 5uM-2 | CA09 | Influenza HA (H1) | NC_026433.1 |
| 5uM-3 | Hawaii2019 | Influenza HA (H1) | MN976432.1 |
| 5uM-4 | MICH15 | Influenza HA (H1) | MK622940.1 |
| 5uM-5 | Brisbane | Influenza HA (H1) | MN569031.1 |
| 5uM-6 | X31 | Influenza HA (H3) | DQ874876.1 |
| 5uM-7 | TX50 | Influenza HA (H3) | KC892952.1 |
| 5uM-8 | SW13 | Influenza HA (H3) | EPI543739 |
| 5uM-9 | SING2016 | Influenza HA (H3) | EPI2053182 |
| 4um-2 | HK2014 | Influenza HA (H3) | AKS48059.1 |
| 4um-3 | NETH1999 | Influenza HA (H2) | CY122252.1 |
| 4um-4 | VN2004 | Influenza HA (H5) | HM006759.1 |
| 4um-5 | CAN04 | Influenza HA (H7) | CY015013.1 |
| 4um-6 | HongKong2013 | Influenza HA (H9) | NAEPI498037 |
| 4um-7 | INF NP | Influenza Nucleocapsid protein | NCAP_I34A1 |
| 4um-8 | INF NS1 | Influenza Non-structural protein | AAA43536 |

### 2.2 Recombinant antigens

#### Production of pre-fusion recombinant Spike (SP) proteins

Ectodomain SP pre-fusion trimers (SARS-CoV-2 S_14-1211_) were produced by co-transfecting plasmids encoding SP-AviTag and SP-6X-HisTag constructs into FreeStyle 293-F Cells at a 1:2 ratio. Transfected cells were cultured in FreeStyle 293 medium for 3 days and recombinant SP trimers were purified from culture supernatant by FPLC using Nickel-affinity chromatography. Purified proteins were biotinylated *in vitro* using BirA enzyme.

#### Production of SP subdomains

NTD (SARS-CoV-2 SP_14-305_) and RBD (SARS-CoV-2 SP_319-541_) monomers with a C-terminal dual AviTag/6X-HisTag sequence were produced by transfecting single plasmid constructs into FreeStyle 293-F Cells. Following a 3-day expression, subdomains were purified from culture supernatant by FPLC using nickel-affinity chromatography and biotinylated by addition of BirA.

#### Production of CoV nucleocapsid protein (N)

Recombinant N containing the full-length N and tandem AviTag/6X-HisTag sequence was produced by co-transforming Rosetta cells with the N expression plasmid and an inducible BirA expression plasmid. Bacterial cultures were grown in the presence of chloramphenicol, ampicillin, and streptomycin, induced with IPTG and supplemented with biotin. Biotinylated N protein was purified by FPLC using a nickel-affinity column and subsequent size exclusion chromatography.

#### Recombinant influenza HA protein production

The nucleic acid sequence corresponding to amino acids 18-524 of influenza HA ectodomains were synthetized (GeneArt, Regensburg, Germany) for influenza virus strains listed in Table 1, as previously described [19]. Synthetic constructs were cloned into the pCXpoly(+) vector modified with a human CD5 leader sequence and a GCN4 isoleucine zipper trimerization domain (GeneArt) followed by either a 6XHIS tag (HA-6XHIS construct) or an AviTag (HA-AviTag construct). The HA–6XHIS and HA–AviTag constructs for each HA were co–transfected using 293Fectin Transfection Reagent into FreeStyle™ 293–F Cells (ThermoFisher Scientific) at a 2:1 ratio. Transfected cells were cultured in FreeStyle 293 expression medium for 3 days. The supernatant was cleared by centrifugation. Recombinant HA molecules were purified from these culture supernatants by FPLC using a HisTrap HP Column (GE Healthcare) and eluted with imidazole gradient. Purified proteins were biotinylated *in vitro* using BirA enzyme.

#### Production of influenza NP and NS1 proteins

The coding sequence of NP from PR8 was synthesized in frame with the coding sequence for a 15–amino acid biotinylation consensus site [20] on the 3′ end (GeneArt). The modified NP sequence was cloned in frame to the 6X-HisTag in the pTRC–His2c expression vector (Invitrogen). NP protein was expressed in the BirA biotin ligase producing *E. coli* strain CVB101 (Avidity), purified by FPLC using a HisTrap HP Column (GE Healthcare) and eluted with a 50–250 mM gradient. The coding sequence for the influenza virus A/PR8/34 NS1 gene (amino acids 3–230, Accession Number: AAA43536) was chemically synthesized (GeneArt) in frame with a 3’ addition of the coding sequence for the BirA enzymatic biotinylation consensus site and the 6X-HisTag. Two mutations, R38A and K41A, demonstrated to prevent aggregation of NS1 at high concentrations, were introduced into the NS1 gene by site-directed mutagenesis. The recombinant NS1 gene was cloned into the pTRC–His2c expression vector (Invitrogen) and expressed in the BirA-enzyme containing *E. coli* strain CVB101 (Avidity). Biotinylated recombinant NS1 was purified by FPLC using a HisTrap HP Column (GE Healthcare), eluted with a 50–250 mM imidazole gradient, buffer exchanged and concentrated.

#### Antibodies and standards

IG bound to microparticles was detected using fluorescent goat polyclonal anti-IG F(ab’)2 secondaries, SouthernBiotech (IgM cat 2022-02, IgG cat# 2062-09, IgA cat# 2052-09). Isotype standards were generated by performing the array on mixtures of IG capture beads with 0.75x serial dilutions of purified human Abs SouthernBiotech (IgG cat# 0150-01, IgM Cat# 0158L-01, IgA cat#0155L-01) ranging from 1µg/mL to 1.3 ng/mL. Anti-SARS Abs JTK137_H22 and JTK137_K6 were generated from the C SP-binding B cells derived from a vaccinated donor. The C SP reactive Ab CR3022 was acquired from IDT Biologika. C SP reactive Ab AHA001 was purchased from Sanyou Bio. HA-reactive Abs (P52d7_K12, P52d7_I1, and P52d7_D20) were generated from HA-binding B cells from a vaccinated donor [21].

Polyclonal anti-C SP reactive Ab was purified from the serum of a vaccinated donor using affinity chromatography. Briefly, 0.5 mg of purified C SP trimers generated as described above was conjugated to cyanogen bromide activated Sepharose (Cytica 17043001) following manufactures protocol. 10mL serum was diluted into 40mL PBS and filtered using a 0.45µm syringe filter (Corning 431220). C SP conjugated Sepharose was added and rotated overnight at 4°C. Resin was collected by centrifugation (2000Xg 10min) and washed twice with 50 mL cold PBS. After the final wash, resin was loaded into a disposable micro column (Pierce 89868) and 0.1M glycine pH 2.8 was used to elute Ab (5 fractions, 200µl) into 25µl 1M tris pH 9.0. Eluted fractions were pooled and concentrated by centrifugation through a 30K centrifugal filter (Millipore Sigma UFC500324). Following concentration, samples were buffer exchanged into PBS using a 0.5mL Zeba desalting column (Fisher scientific PI87767) and quantitated.

### 2.3 Assay protocol

Serum samples were diluted into PBS (1/7150 for IgG detection, or 1/500 for IgM and IgA detection) and 40ul was arrayed in 96 well u-bottom plates. A 5µl suspension containing a mixture of each antigen-coated microparticles (5e5 beads per antigen). The suspensions were mixed by pipetting and incubated for 15min RT. The beads were washed with 200µl PBS and centrifuged (3000Xg 5min RT). The CBA particles were resuspended in a secondary staining solution consisting of the appropriate anti-Ig secondary diluted 1/400 in 1%BSA containing PBS. The suspension was incubated for 15min RT in the dark. The beads were washed with 200µl PBS and pelleted (3000Xg, 5min RT). The particles were resuspended in 80µl PBS and analyzed on a BD Cytoflex flow cytometer in plate mode at sample rate of 100µl/min. Data collection was stopped following acquisition of 75 µL of sample. Following acquisition, the resulting FCS files were processed and analyzed.

### 2.4 Automated analysis and software

We developed the CBA Toolbox to automatically process FCS files to rapidly quantify the antigen binding by serum polyclonal Abs or purified Ab samples. This software was developed in Matlab (The Mathworks, Inc. Natick MA, USA) version R2020a on MacOS. It requires the Statistics and Machine Learning Toolbox, the Curve Fitting Toolbox and the Signal Processing Toolbox. Additionally, code from Matlab Central (www.mathworks.com/matlabcentral/) public code repository was incorporated into this program [22–25]. This software is available on GitHub [https://github.com/UAB-Immunology-Institute/cba-toolbox].

### 2.5 Human serum samples

Human serum samples were collected from de-identified deceased organ donors. Peripheral blood from human subjects was drawn into K2-EDTA tubes (BD Bioscience). Plasma was isolated by density gradient centrifugation over Lymphocyte Separation Medium (CellGro), aliquoted, and stored in LN_2_. Cryopreserved samples were thawed at RT and diluted 1/100 in PBS in a 96 well U bottom plate to generate a ‘master plate’ stored at −20°C. Final dilutions were generated in assay plates as needed. For initial experiments focused on assay optimization and serum dilution determination, samples were acquired from consented participants under the UAB Institutional Review Board approved the study protocol (IRB-300005127).

## 3. Results and discussion

### 3.1 Generation of antigen loaded microparticles

To evaluate the possibility of using antigen loaded beads to quantitate specific Ig, we adsorbed CA09 HA and C SP onto two distinct SA conjugated particles and stained each with the I1 HA reactive and CR3022 C SP-reactive mAbs. The MFI of binding by these two mAbs to their cognate antigen coated particles were nearly identical when tested at 100µg/mL to 1ng/mL concentrations (FIG 1a) demonstrating that the linear range of the CBA array was nearly three orders of magnitude and that the assay could reproducibly detect specific Abs at a concentration as low as 10ng/mL Additionally, the MFI curves as a function of concentration were also nearly identical, suggesting that when antigen loading is carefully controlled to be equivalent, the MFI signal is dependent on mAb concentration irrespective of the antigen.

**Figure 1.**
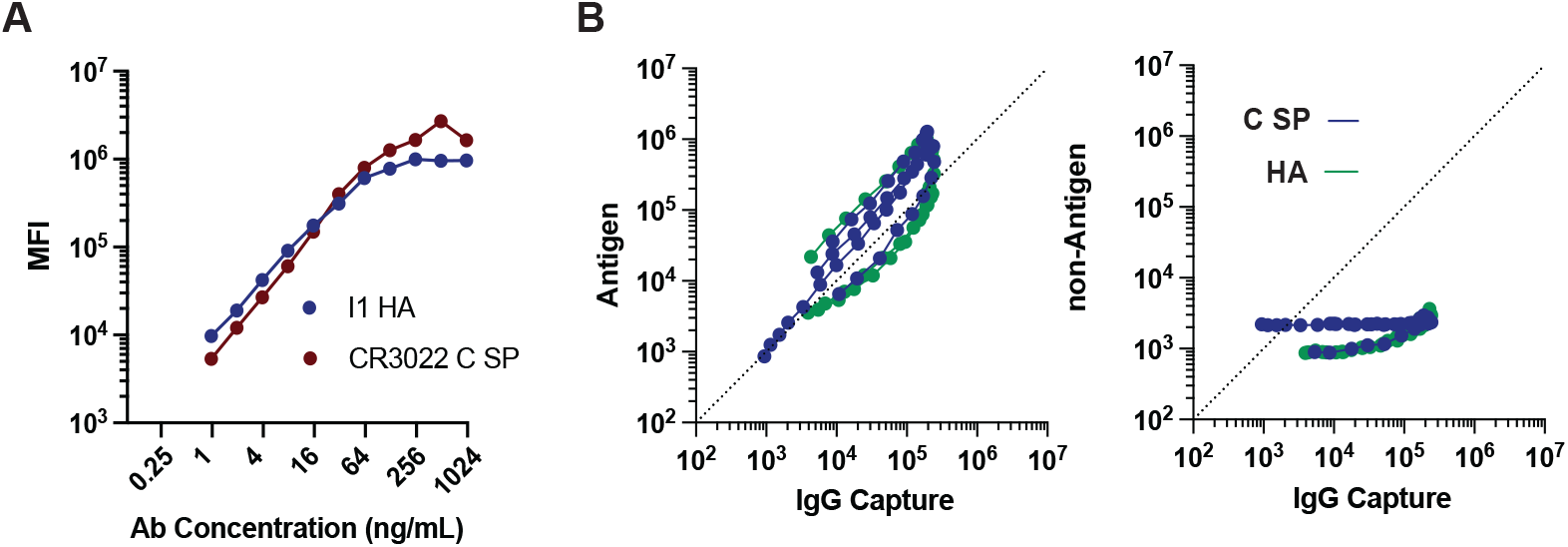
Dilution curves of antigen-specific antibodies. **A.** Plots of the MFI of beads coated with either CA09 HA or C SP and as a function of the concentration of the indicated mAb. **B**. Plots of the MFI of IgG capture beads and HA (green) or C SP (blue) coated beads simultaneously stained with dilutions of HA or C SP reactive Abs. Curves from the cognate antigen (left panel) or non-cognate antigen (right panel) are shown.

Because the derived MFI represents the association of the fluorescently labeled secondary Abs with the bead:Ag:Ab complex it is independent of the nature of the antigen, we therefore considering using beads coated with anti-Ig to allow quantification of bead bound Ab. We generated IgG capture beads by absorbing biotinylated anti-human IgG Fab(2) to the same lot of SA conjugated beads coated with HA and C SP. We then incubated these reagents with additional mAbs specific for either C SP (CR3022, AHA001, H22, and K6) or HA (K12, I1, and D20) and compared the MFIs following staining with fluorescently conjugated anti-IgG secondaries.

As seen in Figure 1b, the MFI signals derived from both the antigen coated and Ig capture conjugated beads were highly similar across the dilution range toward their cognate antigen and exhibited little or no binding to their non-cognate beads. The MFI Ag/Ig bead ratio differed slightly among mAbs likely reflected differences in the affinity of these mAbs for their cognate antigen, as has been described for these HA reactive Abs [21]. These data confirm that biotinylated anti-isotype capture Abs, when the antigen density and capture Abs on the beads was equivalent, mirror the quantity of antigen bound Ig and serve as an indirect standard for antigen reactive Ab quantification.

### 3.1 Generation of the β-COV and INF arrays

To simultaneously interrogate Ab reactivity to multiple viral protein antigens, we expanded this array to include all particles available in the 4um and 5um kits which have distinct inherent fluorescence in the far-red channels (FIG 2ab). We maximized SA conjugation such that the binding capacity of each bead was high and uniform. To ensure equivalent SA loading across bead sets, the biotin binding capacity was evaluated by incubating SA bead preparations with an Alex-488 labeled recombinant influenza A/PR/8/34 hemagglutinin (PR8 HA) trimer that was biotinylated at a single site on the C-terminus of the trimer (FIG 2b). As expected, the intensity of bead labeling by the fluorescent HA trimer was dependent on the amount of HA protein in the reaction. Not surprisingly, the 4µm and 5µm particles exhibited different maximum intensities due to the difference of surface area (therefore differences in total SA bound) between the two particles sizes (FIG 1bc). However, SA loading was uniform when comparing particles of the same size, with a coefficient of variation of 5.39% and 5.34% for all 5µm and 4µm beads, respectively.

**Figure 2.**
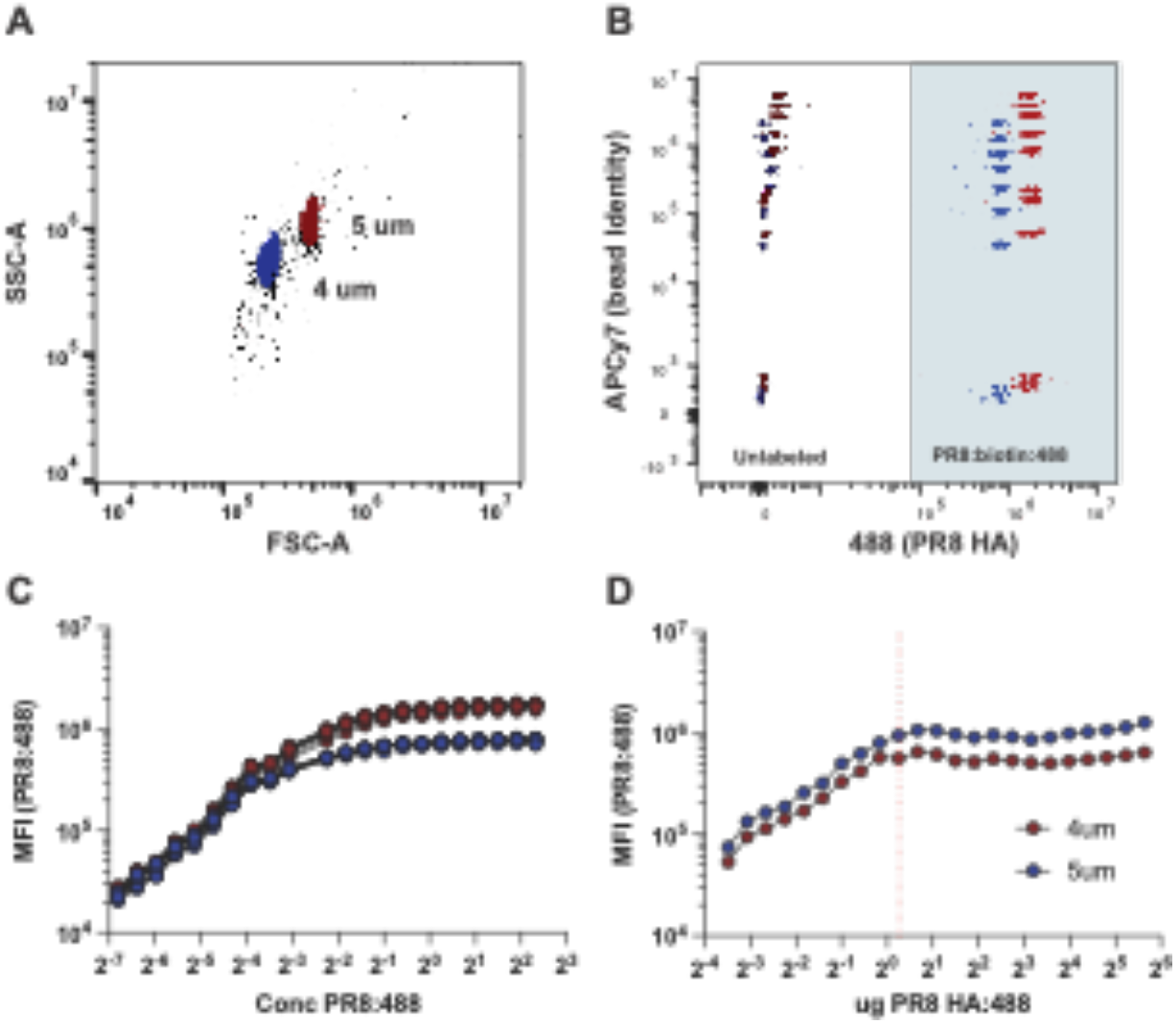
Array generation and loading controls. **A.** Forward and side scatter histograms for differentiating the indicated size particle. **B.** Flow cytometric histogram of the array constituents before (left, unshaded) and following incubation with biotinylated / alexa488 PR8 HA (right, shaded). **C.** MFI of each array constituent stained with the indicated concentration of PR8 HA:biotin:488. **D.** MFI of the indicated beads stained with increasing amount of PR8 HA:biotin:488 per 1e5 particles. Amount at which beads become saturated is indicated by the vertical red dashed line.

Next, we determined the biotin binding capacity of the SA-conjugated beads. First, we incubated the SA-conjugated particles with an excess of a known amount of biotinylated influenza HA trimers and measured the protein absorbed from these suspensions using UV spectrometry. For the two independent lots of SA-conjugated beads we tested, an average equivalent of 220µg HA protein per 1X10^8^ particles was absorbed. Second, we incubated the SA beads with decreasing concentrations of fluorescently labeled PR8 HA and determined the mean fluorescence intensity (MFI) of binding (Fig. 2D). The point of inflection, at which the MFI of the beads progressively diminished as a function of decreasing PR8 HA concentration indicating a sub-saturating quantity of biotin, equated to the same value as determined by the first method – specifically 200µg of PR8 HA per 1e8 beads. Based on these two approaches, we estimated the typical biotin binding capacity of the SA beads to be 0.1 picomole biotinylated protein per SA conjugated particle.

To generate antigen arrays, we absorbed recombinant viral proteins to the individual fluorescent microparticles (Table 1). β-CoV array consist of full length, pre-fusion stabilized SP trimers derived from the 5 known β-CoV: SARS-CoV-1 (S), SARS-CoV-2 (C), MERS (M), HKU1 (H) and OC43 (O). In addition to the monomeric Receptor binding domain (RBD) subunit of SP from SARS-CoV-1 and SARS-CoV-2 and the SP N-Terminal Domain (NTD) from SARS-CoV-2. Finally, INF HA (CA09 HA) was included as a positive control in the array as Abs specific for recently circulating INF strains or vaccine constituents can be found in almost all adults.

For the INF array, we absorbed HA derived from 14 influenza viruses to microparticles listed in Table 1b, in addition to INF NP and NS1. These HA represented the most common circulating H1 and H3 strains over the last decade, as well as PR8, a common laboratory strain, and a member of H2, H5, and H7 strains.

### 3.4 Antibody standards and quantification

Although the data above suggested that the CBA could provide sensitive qualitative assessment of antigen-specific Ab levels, we sought to refine the assay to accurately measure the amount of antigen-specific Ab present in biological specimens. We postulated that the relationship between Ab concentration and MFI derived from the CBA analysis of these Abs would be dependent on several factors, including the Ab isotype, which is unequally recognized by pan IgG secondaries, the affinity of the Ab for the antigen, and the valency of Ab-antigen interactions. To normalize minor deviations in biotin binding capacity between the individual array components, and the notable differences between particle sizes, and we incorporated all individual particles into a panel of Ig-capture beads distributing anti-IgM, -IgA, and -IgG capture beads across all the beads used for the antigen arrays (FIG S1a). In parallel to the antigen specific array, Ig capture bead mixtures were stained with purified IgM, IgA, or IgG serially diluted (0.75x) from 1ug/mL to ∼1ng/mL. The resulting standard curves were then used to extrapolate the Ab isotype bound to the antigen-specific beads thus providing an accurate quantification (FIG S1b).

### 3.3 Assay functional range and dilution factor determination

CBAs, just like conventional ELISAs, can exhibit prozone effects when the concentration of the Abs being evaluated exceeds the binding capacity of the array. Although prozone effects are less prominent in assays with a larger dynamic range, it is still important to choose analyte dilutions that allow for accurate extrapolation of the MFI measurements to concentration. This is particularly important for polyclonal sera analyses. To identify a dilution range that would facilitate accurate measurements of viral antigen specific IgG, IgM, and IgA serum Abs, we diluted (between 1/500 and 1/20517) serum samples from ten SARS-CoV-2 convalescent patients and analyzed the samples using the β-CoV CBA. As shown in Figure 3, the maximum MFI of binding by C-SP reactive IgG Abs to the C-SP coated beads was significantly higher when compared to maximum binding by C SP-specific IgA and IgM Abs. For samples that contained detectable anti-C SP IgG, the prozone effect was apparent at dilutions below 1/7000, after which the signal decayed as a function of Ab concentration. By contrast, IgM and IgA Abs specific for C SP, which could be detected at a 1/7000 dilution in some samples, did not exhibit a prozone effect at this dilution. This difference was likely due to the relatively lower concentrations of C SP-specific IgA and IgM Abs present in serum [26] or to differences in signal intensity, which is influenced by the isotype-specific secondaries. Using this experiment as a guide, we determined that a 1/7100 dilution of convalescent SARS-CoV-2 serum was appropriate to measure C SP-specific IgG while a 1/500 serum dilution was appropriate to measure C SP-specific IgM/IgA. However, the appropriate dilution of serum necessary to avoid prozone effects in the CBA must be empirically tested and is likely to be dependent on the Ab levels within the experimental samples being assessed and the antigen of interest.

**Figure 3.**
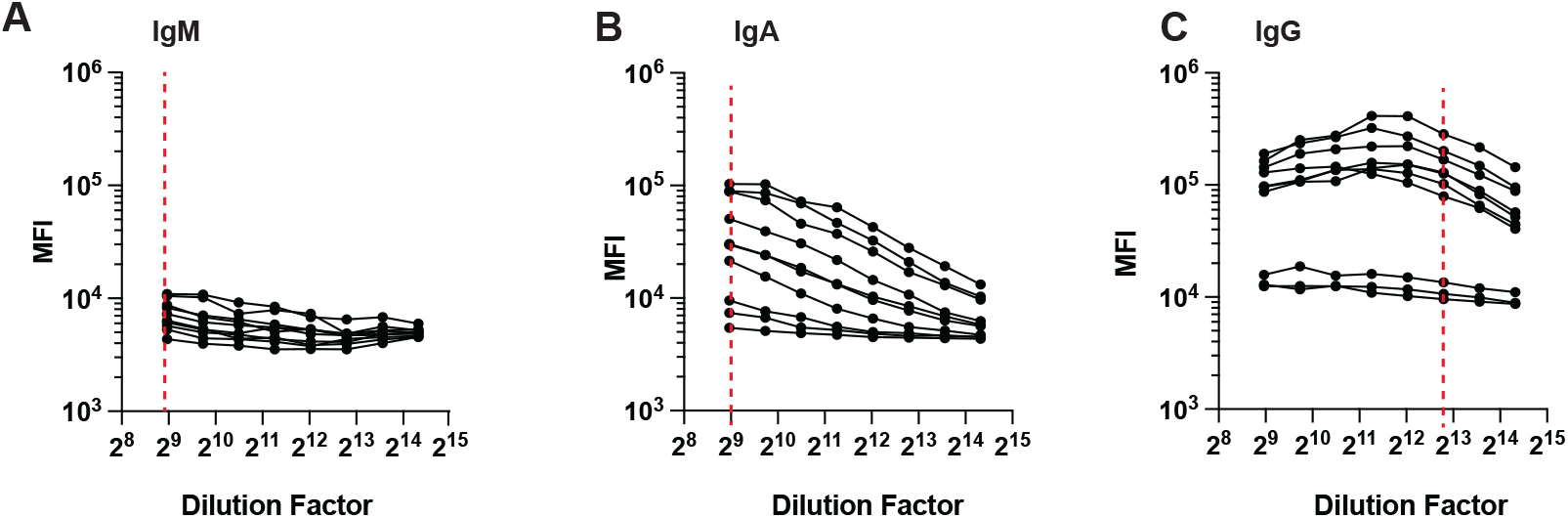
Optimal assay serum dilution determination. MFI of CS SP Beads stained with serial dilutions of human serum from COVID convalescent patients visualized with secondaries reactive to the indicated human Ig isotype. Serum Dilutions selected to assay for each isotype are indicated by red dashed lines.

### 3.5 Automated Analytic Workflow

Although the array data, which was collected as FSC files using a traditional flow cytometer, could be analyzed using many different flow cytometric software packages, the inclusion of multiple standard curves and the large number of individual flow files made manual data processing laborious and subjective. To circumvent these issues we developed a software pipeline to perform automated sample quality assessments and batch processing. This software, written in Matlab (The Mathworks, Inc., Natick MA, USA), processed individual raw FCS files to identify events associated with the beads, to find bead peaks and to summarize the raw data. The software analyzed the standard controls and automatically back-calculated concentration units from raw MFI data. The software was also designed to work in batch mode, which allowed for processing many files at once.

The overall workflow is shown in Figure 4. The command line program required four arguments: (i) a directory containing all the FCS files; (ii) a sample manifest containing metadata for each of the corresponding FCS files, including the flow cytometry channel mappings for the bead channel and the secondary isotype channel; (iii) a configuration file describing the association of bead peak order (numbered from lowest to highest FL4 MFI) with protein for the specific assay, (iv) and a configuration file containing parameters for peak identification. The output was a Matlab structure array where each element contains a table of isotype reactivity for each feature (individual bead/isotype MFI) in the assay as well as other program settings for complete analytical provenance supporting reproducibility.

**Figure 4.**
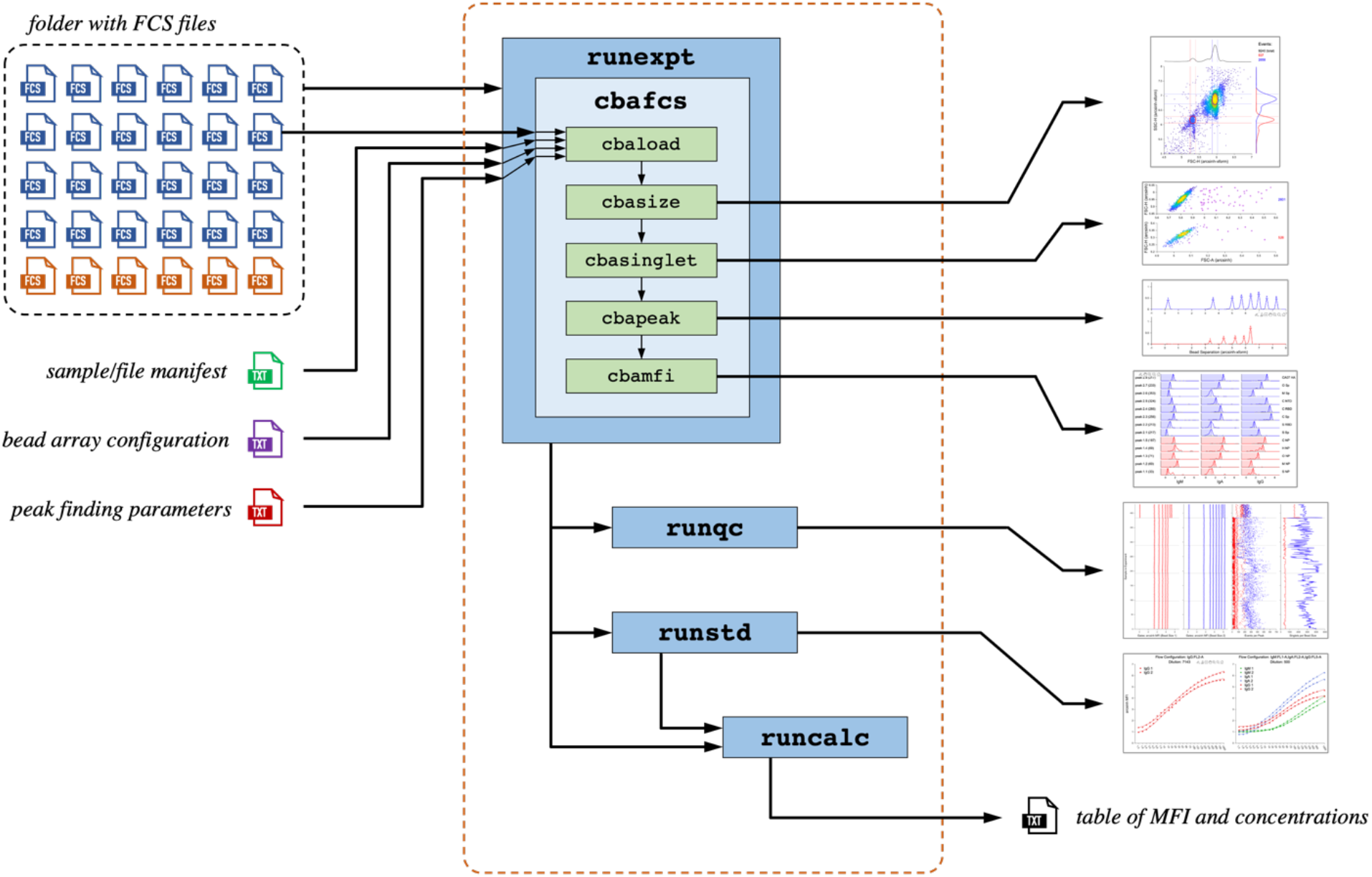
Schematic of automated software modules. Software components are shown within red dashed box. Green boxes are modules that perform a single element of processing. The light blue box is a program that processes a single FCS file to extract transformed MFI data for all features in the assay. The darker blue box represents a wrapper to batch process many FCS files (represented in black dashed box) and annotates whether an FCS file is a normal sample (blue) or a standard control sample (orange). A sample manifest is used in batch mode to provide sample metadata. Additionally, a bead array configuration file and a peak finding parameter file contain parameters that describe association of beads with proteins and settings applied to each file. After the program is run in batch mode, there are modules for bead gate quality assessment, construction of standard curves and back-calculation to concentration units. When running for an individual file, figures can be generated for diagnostic purposes (right).

This structure array, in turn was used for an analysis quality assessment program to visually summarize the consistency of the bead gates for each bead size as well as the numbers of beads for each bead gate as shown in Figure S2. This structure array was also used to generate standard curves for each isotype, bead size and sample dilution (multiple dilutions can be run together and are annotated in the sample manifest, and each requires their own set of standard curve samples). The samples used to generate the standard curves were annotated in the sample manifest. These data were used to compute a four-parameter logistic (4PL) fit for each bead size/isotype/dilution as shown in Figure S1b Finally, the structure array and the 4PL fits were used to back-calculate concentration units for the hyperbolic arcsine-transformed MFI data. The software program then outputs the entire data set to a single, tabular text file that was suitable for importing into any analysis program or relational database for large-scale studies. This software was also suitable for high-throughput use as data derived from five 96-well plates run on a Cytoflex (Beckman Coulter, Indianapolis IN, USA) – 480 samples including standards – could be processed in about a minute on a current model Macbook Pro laptop with no special memory or processor requirements.

### 3.6 Deriving MFI Reactivity Data from a Sample

The computational workflow was designed to parallel the process for manually quantifying the concentration of antigen-specific Abs in a sample (Figure 5). First, the MFI datapoints for each sample were extracted from the FCS files [22], transformed using a hyperbolic arcsine [23] and then visualized in a forward-scatter vs. side-scatter plot. The beads of different sizes (4um and 5um) were automatically detected by looking for the expected number of peaks along the forward-scatter (x) axis to define gates, and then, within each gate, to identify single peaks on the side-scatter (y) axis. The intersection of these gates in 2-D space was used to define oval gates (FIG 5a). Events that were present in each oval gate were annotated as distinct bead populations. Next, forward-scatter height vs. area plots for events in each bead gate were evaluated using density-based clustering and outliers, which represented non-singlet beads were then excluded (FIG 5b). Singlet beads were then used for peak finding (for each bead size gate) based on transformed MFI of the specified bead channel specified in the manifest header and compared against the expected number of peaks as indicated in the array configuration file (FIG 5c). Finally, events from each bead gate were evaluated on the secondary isotype flow channel(s) specified in the manifest header to quantify the location of the peak (in transformed MFI) for each bead feature and isotype (FIG 5d).

**Figure 5.**
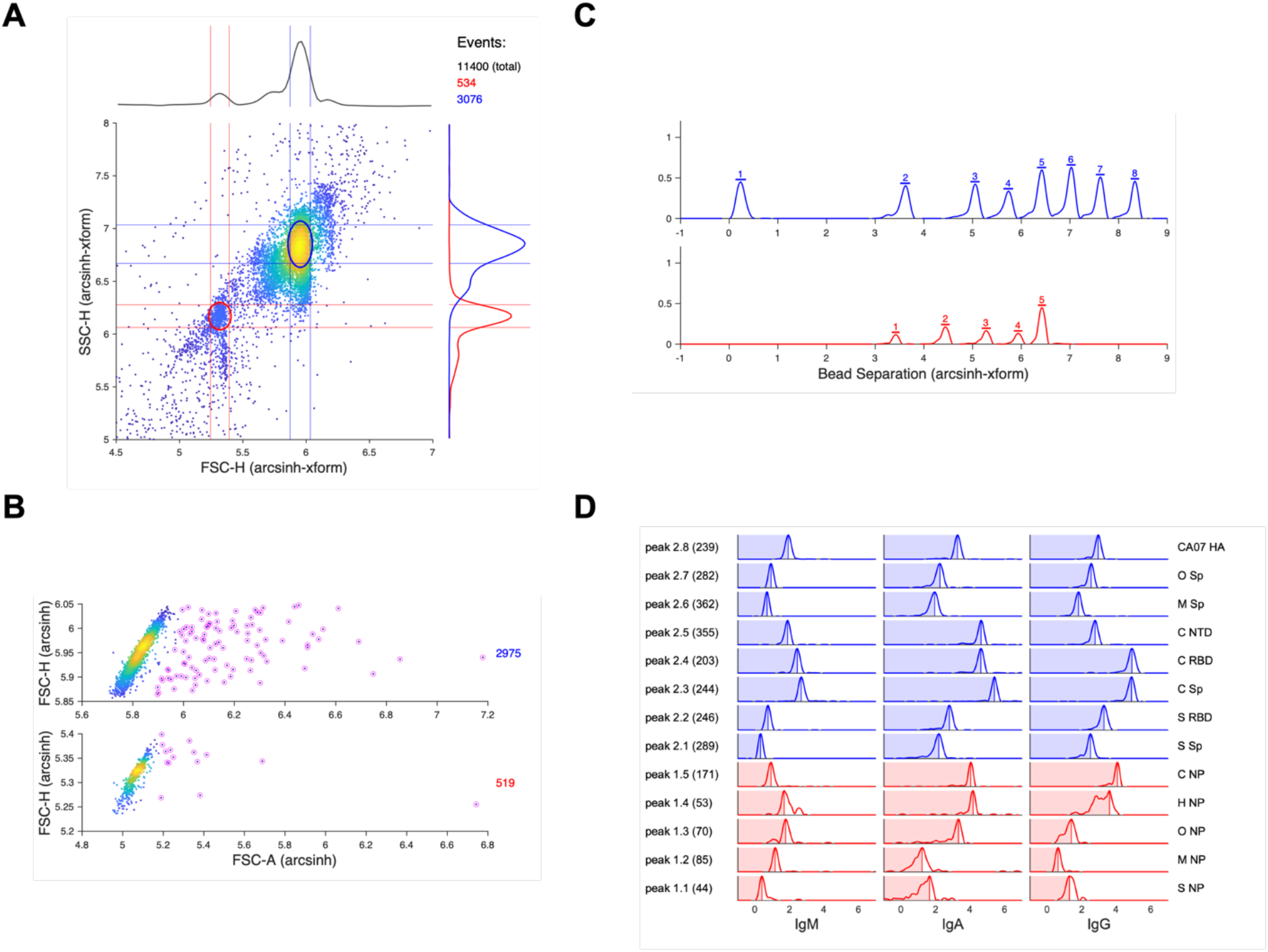
Raw FCS data processing steps. All intensity data is hyperbolic arcsin transformed. A. Forward scatter vs. side scatter to identify densities of events associated with multiple bead sizes. Red and blue ovals are gates for 4um and 5um beads, respectively. B. Density-based clustering on forward scatter height vs. area to identify outliers, or non-singlets for each bead size gate. C. Identification of peaks on the specified bead channel for each bead size after non-singlets filtered out. Line segments above each peak are individual bead gates. D. Hyperbolic arcsine MFI intensity plots on the secondary isotype channels (columns of plots) for each bead size/peak gate (rows of plots). For each row, number of beads per gate indicated in parentheses. Antigen features for this example panel are: C = SARS-CoV-2; S = SARS-CoV-1; M = MERS; O = OC43; H = HKU-1; CA07 HA = California 07 influenza hemagglutinin; Sp = spike protein; RBD = receptor binding domain; NP = nuclear protein; NTD = n-terminal domain.

This software can accommodate an optional “split channel” prior to bead peak detection (Figure 5c). By labeling half the beads prior to flow and employing an otherwise unused flow channel to detect this signal, the number of antigens profiled can be potentially doubled. Importantly, comparison of MFIs computed with the automated software to manual processing using FlowJo (BD Biosciences, Franklin Lakes NJ, USA) led to similar results, as shown in Figure S2 for four representative samples.

### 3.7 Polyvalency and subunit corrections

When we began to analyze serum samples and polyclonal Ab preparations, we noted that the highest MFI values observed often eclipsed that recorded using antigen-specific mAbs, suggesting that the use of antigen-specific mAbs standards would result in an overextrapolation of the total antigen-specific Ab in more complex samples, and that the cytometric array is sensitive to effects resulting from the polyvalency of Abs present in sera or other polyclonal Ab preparations. We hypothesized that multiple epitopes present on antigens were simultaneously bound resulting in higher maximum MFI values than exhibited by an equivalent concentration of a mAb, a phenomenon that would influence Ab quantification. To determine if this were the case, and to quantitate this polyvalency effect, we assayed a polyclonal C SP reactive Ab preparation derived from the serum of a vaccinated donor. Using antigen-specific capture beads to quantitate the Ab isotype in this preparation, we determine it was composed of 90 percent IgG, 10 percent IgA, with low but detectable levels of IgM. Because IgA was present as a significant portion C SP reactive Ig, we further purified IgG using protein G chromatography yielding an anti-C SP pIgG with a mixture of IgG subclasses. We then evaluated both the pAb and IgG anti-C SP preparations in the CBA. As can be seen in Figure 6a, the MFI of the IgG detection Abs was noticeably but marginally higher in the IgG fraction resulting from the presence of IgM and IgA Abs in the Ig preparation. We then used the array to quantitate C SP Abs and compared these calculated values to the known concentrations. Although the calculated values were higher than the known concentrations, they exhibited a linear relationship up to concentrations of approximately 0.75 ug/mL, a concentration in which the prozone effects of anti-SP Abs become apparent (FIG 6a). By comparing the known IgG concentration with the calculated values along the serial dilutions within the linear range we could determine the increase in apparent concentration resulting from the polyclonality of the purified anti-C SP Ab, which was an apparent 3.16-fold inflation of the calculated concentration (FIG 6b). These data demonstrate the average polyvalency effect within the assay is constant within the range of concentrations indicated and provided a correction factor for extrapolated Ig concentration values. To further confirm this normalization was accurate, we assayed the NIBSC anti-C SP standards at serial dilutions ranging from 1/1e3 to 1/8.65e4. When the resulting anti-C SP values are corrected and plotted against the known dilutions of the pAb anti-C SP they collapse to near identity, confirming the accuracy of this correction (FIG 6c). Using this approach, we calculate the total anti-SP IgG to be 221.7µg/mL and 90.4µg/mL in the NISBC standards 2-/162 and 21/324 respectively (Fig 6d).

**Figure 6.**
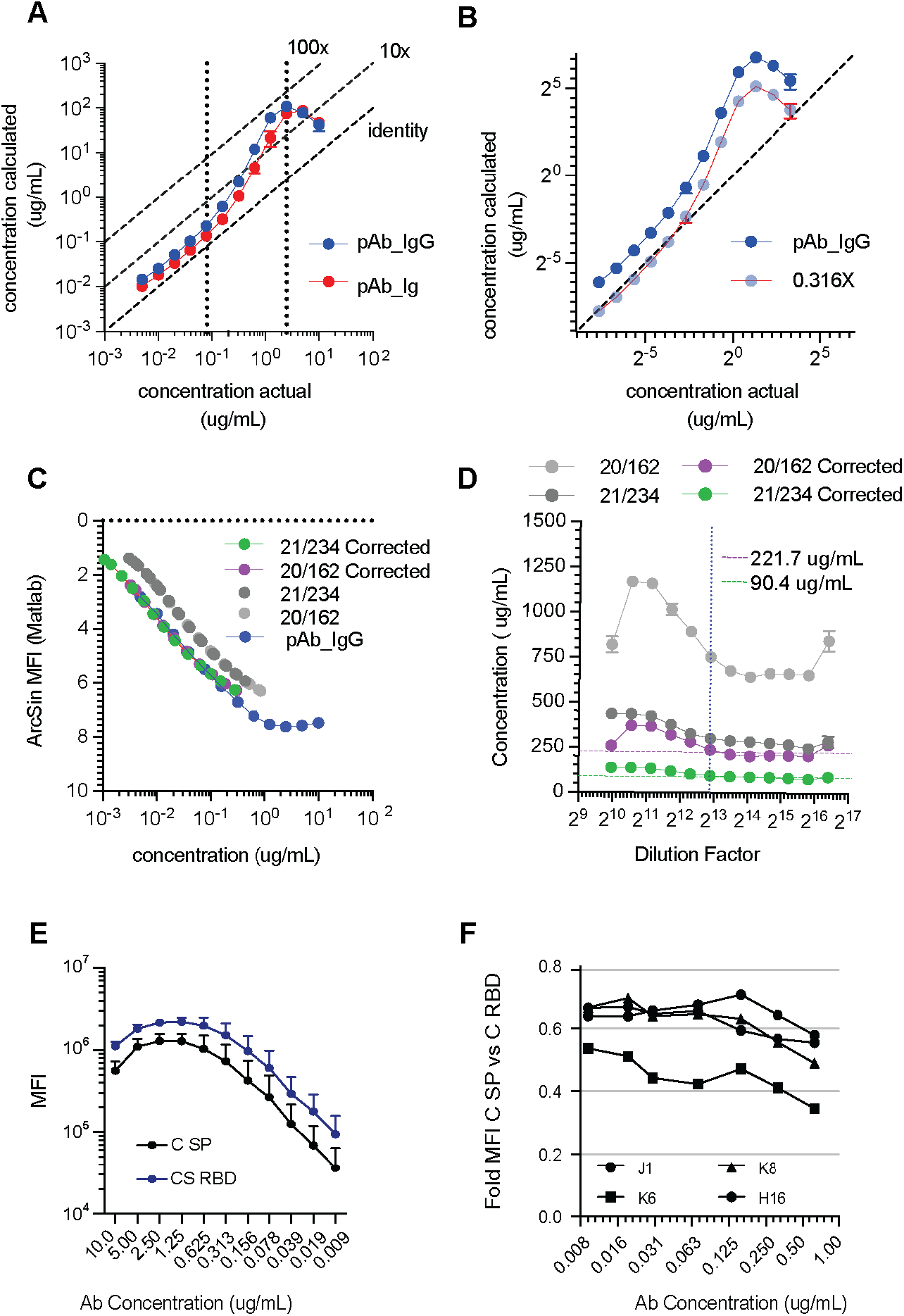
Quantitated polyvalency and C SP subunit corrections. **A.** Extrapolated concentration of anti-CS SP pAb and pIgG plotted against the known concentration. **B**. Calculated polyvalency correction factor for anti-CS SP pAb IgG. Fold difference in MFI between CS SP and CS RBD across the linear range of dilutions is shown. **C**. Hyperbolic arcsine transformed MFI values derived from the array plotted against known concentration of anti-CS SP pAb and uncorrected values derived from the indicated NISBC standards. Corrected values were generated by linear scaling of the extrapolated concentrations. **D.** Extrapolated anti-CS SP antibody concentrations (before and after polyvalent correction) of the indicated NISBC standards. Dashed lines indicated the high confidence Ab concentration of the indicated standards. **E.** Mean MFI of the CS SP coated (black) or C RBD coated (blue) beads stained the indicated concentrations of anti-CS RBD reactive rAbs. The resulting mean and SD of the MFIs of CS SP and CS RBD are shown. **F.** The ratio of the MFI of CS SP to CS RBD of the indicated anti-CS RBD rAbs at the indicated concentration.

In addition to the polyvalency effects described above, we noted that the MFI values were higher for the C RBD particles than for full-length SP trimers. This effect was manifested as exceptionally high extrapolated anti-C RBD values in some samples. Comparing the MFI of the C RBD and C SP beads resulting from staining with dilutions of four RBD reactive mAbs, we consistently observed higher values for the RBD particles (FIG 6e). As these Abs recognize an identical epitope on both antigens and the bead constituents exhibit identical biotin binding capacity, we reasoned that because the RBD is expressed as a monomer and has variable accessibility in the full-length trimers (RBD up-down configurations [27]) the epitope density is unequal between these configurations. To determine if this was a consistent effect, we analyzed the fold difference in MFI between C SP and C RBD beads at a range of concentrations of four anti-RBD Abs (FIG 6f). Although the CBA signal derived from mAbs is dependent on Ab affinity in addition to concentration, three of these Abs exhibit a concentration independent 1.5-fold increase in C RBD MFI compared to that of C SP trimers, allowing us to correct for differences in RBD epitope availability between full length SP and monomers.

### 3.8 Sensitivity, accuracy, and reproducibility

Although the array results are highly reproducible when assaying monoclonal antigen-specific Abs or isotype-specific Ab standards, we sought to determine the variance of array across repeated measurements. To quantitate the reproducibility of the array derived Ab values, we analyzed serum available through the UAB Comprehensive Center for Human Immunology (CCHI) donor repository twice, independently, with COVID and INF arrays. The MFI of each antigen in the arrays was nearly identical in the between the two replicates (FIG 7a). Because acquisition for most of these samples pre-dated the COVID19 pandemic, they did not contain detectable SARS-CoV-2 antigen reactivity (FIG S3). However, IgG reactivity toward one or more HA antigens was observed in all samples. The calculated value of the most prominently targeted HA antigens (including both H1 and H3) were in close agreement between the replicates (FIG 7b). Although deviations between replicates were noted at the highest Ab levels detected, these fall outside of the linear range of the assay. Within the range of confidence, we observed a high level of agreement between the two measurements. Importantly, as shown in Figure 7c, the calculated values of CA09 HA, which was included in the COVID array as a positive control, were nearly identical in both arrays. As these reagents were generated independently using different lots of SA beads, this demonstrates the reproducibility of both bead generation and assay measurements.

**Figure 7.**
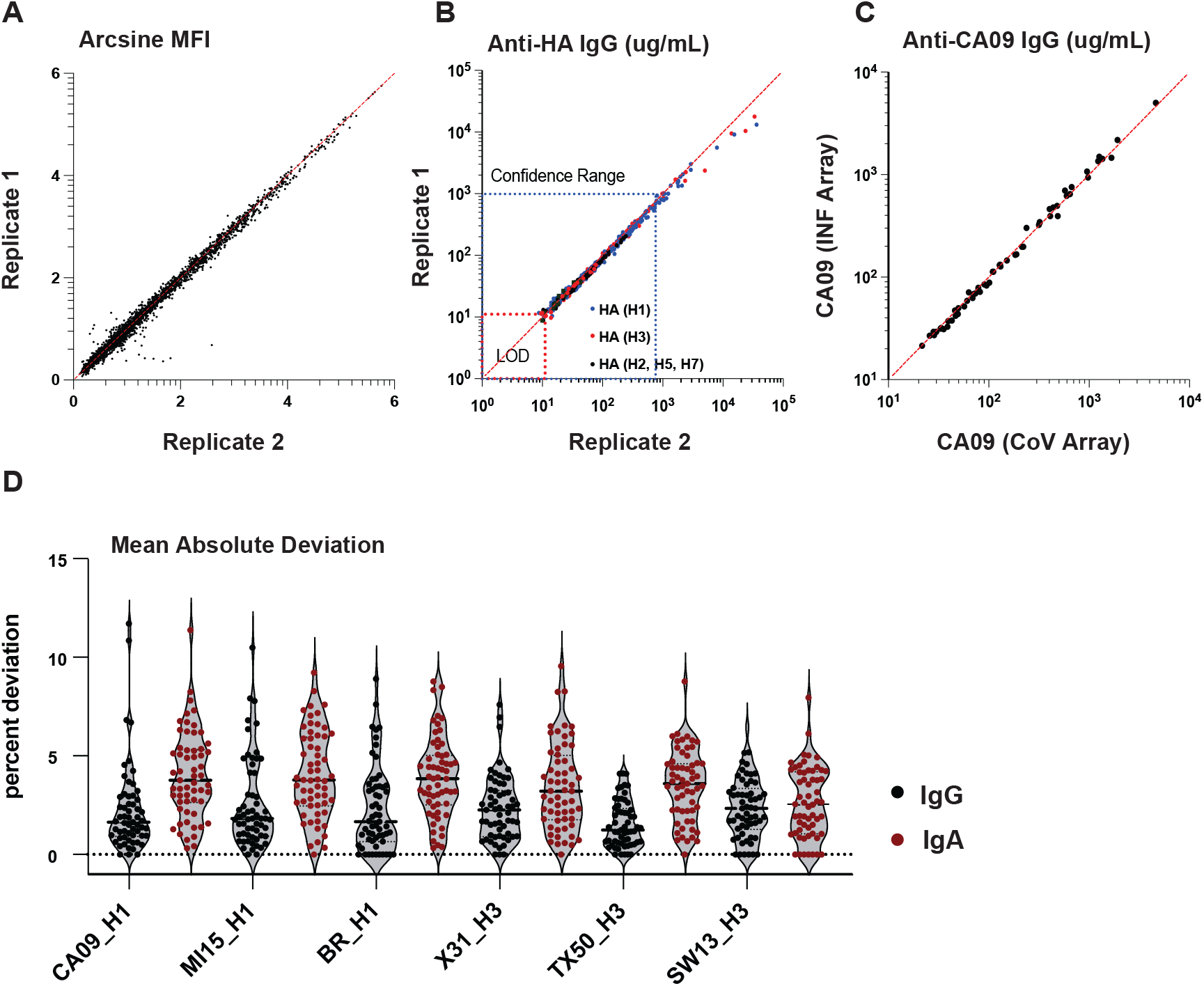
Assay reproducibility. **A.** Scatter plot of the ArcSin MFI of each bead for all samples for two independent measurements of the CCHI U19 cohort serum. The values of the two replicates are plotted pairwise. **B**. Extrapolated antibody concentration for the two replicates ploted pairwise for the indicated HA antigens. **C**. Scatter plot of the derived concentration of Ab reactive with CA09 HA derived from the COVID and INF arrays. **D**. Mean absolute deviation for each data point of two replicates across the two replicates for the indicated antigens and Ab isotypes.

To quantitate the reproducibility of the assay, we examined the mean absolute deviation of IgG and IgA reactivity toward the three most frequently targeted HA H1 and H3 antigens (FIG 7d). The average deviation is less than 5% across all these antigens, and slightly lower for IgG (2.5%) than for IgA (4%). Collectively these data suggest the arrays are highly reproducible both at the level of bead generation and in their performance.

We did not observe Ab binding to C SP coated beads in serum samples collected before the spring of 2021, despite IgG reactive with O SP being readily quantifiable in all samples (serum antigen-specific Ab concentrations for all antigens is shown in Figure 8). Following the spring of 2021, C SP reactivity was observed in 13/19 samples whereas C NP reactivity was only observed in 4. In every case, C NP reactivity was corelated with C SP Ab, suggestive of a prior infection (FIG S3). These data are consistent with the infection and vaccination rates within the general population from which these donors were derived [28].

**Figure 8.**
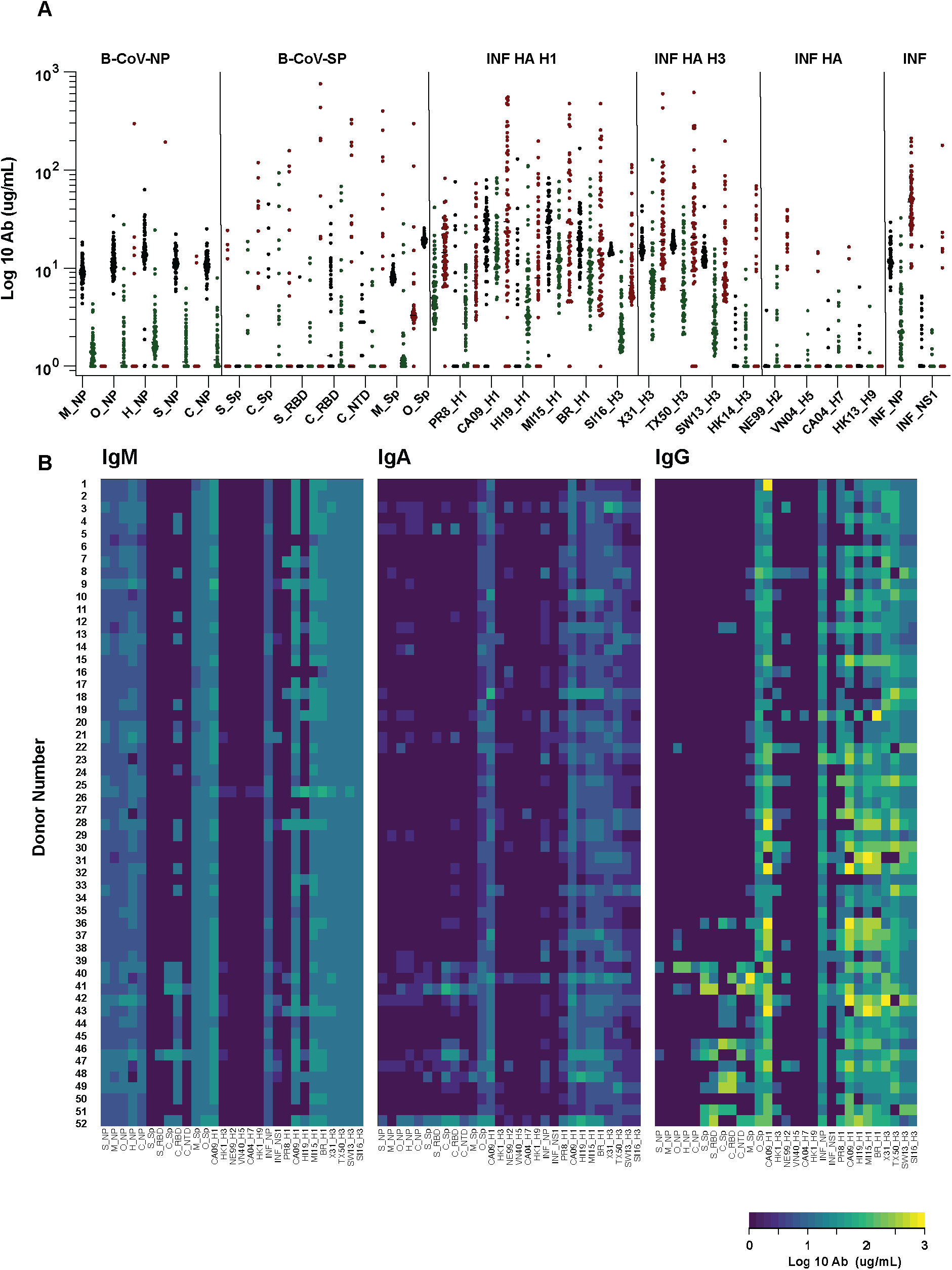
The reactivity profiles of 96 sera samples toward the full β-Cov and INF arrays. **A.** The concentration of IgM (black), IgA (green), and IgG (red) reactive with the indicated antigen as measured by the Coba and INF array. **B.** Heatmap of the log10 Ab concentration (ug/mL) for the indicated antigen. Each row corresponds to an individual serum, and each column corresponds to the indicated antigen, IgM, IgA, and IgG are shown.

Although HA reactivity was observed in all samples, there were significant differences in the HA reactivity and Ab levels observed across individuals, representative array histograms shown in Figure S4. A bias toward either H1 or H3 HA was observed in some individuals, while others exhibited a balanced anti-HA Ab repertoire, although the recent vaccine formulations were apparent, with CA09, Brisbane (H1) and MI15 demonstrated the highest level of reactivity. Reactivity toward PR8 was observed to be the lowest of H1 HA (FIG 8). Reactivity toward H3 were generally lower than that of H1 while reactivity toward the H2, H5, H7 and H9 included in the array were rarely detected. Most individuals exhibited IgG and IgM Abs toward INF NP while only 10% harbored serum reactivity toward INF NS1, perhaps indicative of a recent INF infection. When we compared the reactivity of these antigens, we noted a significant correlation between H1 and H3 HA, which were poorly correlated with one another (FIG 9a). Although we cannot determine if this is due to cross-reactive Abs, or that similar levels of Abs that target these antigens, high sequence similarity, as is the case for CA09 and MI15, are associated with the highest correlation of Ab levels (FIG 9bc). Interestingly, although IgG and IgA Ab levels are both highly corelated toward similar HA antigens, the IgG and IgA levels themselves do not correlate (FIG 9cd), perhaps suggestive of different origins of HA reactive Abs of these isotypes. Collectively these data demonstrate the assay is suitable for high-dimensional, high-throughput Ab reactivity determination.

**Figure 9.**
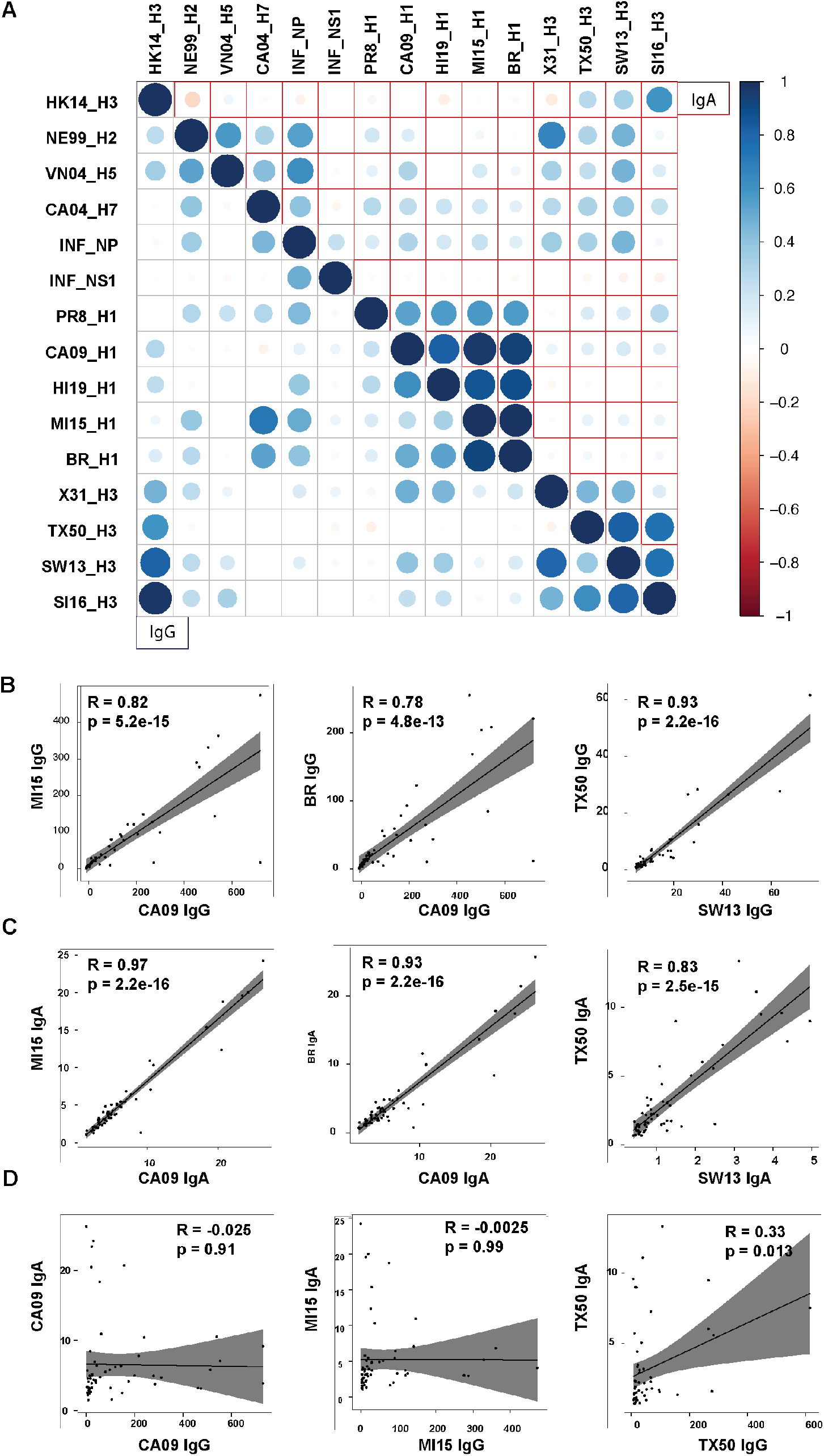
Correlation of Ab levels for HA antigens in the array. **A.** The pairwise Pearson correlation coefficient for either IgG (lower left) or IgA (upper right) of Ab levels for each antigen in the INF Array. **B.** Scatter profiles of antibody levels toward the indicated H1 and H3 hemagglutinins for the 59 samples in the cohort.

## 4. Conclusions

Here, we describe an CBA platform designed to conduct high-throughput determination of Abs reactive with viral antigens. The assay provides a rich view of the humoral immune landscape by multiplexing the analysis of 12 β-CoV or 16 INF antigens into a single assay. By altering dilutions factors and secondary detection reagents, the levels of IgG, IgM, and IgA can be accurately determined. This platform exhibits high sensitivity, although the lower limit of detection is dependent on dilution factor, as little as 2ng/mL of antigen-specific IgG in the final dilution can be measured. The assay demonstrates high specificity, as demonstrated by multiple monoclonal Abs, the reactivity to each antigen in the array is discrete. The use of multiple indirect standards provides accurate quantification for Ab reactive to each antigen across all assayed isotypes. Additionally, the assay requires minute amounts of both antigen and serum tested, typically consuming less than 0.5µg of antigen and less than 1µL of sample per assay.

Although, the CBA array described here is focused on β-CoV and INF serology, the same strategy can be implemented to evaluate serum Ab to a wide array of antigenic targets. The use of recombinant antigens, like the various SP ectodomains and subunits or HA and NP described here, allow for the rapid replacement or addition of similar antigens into the CBA with no required modification to the analytics. Although the corrections for polyvalency and subunit epitope availability are specific for C SP, these are the product of the physical nature of the antigens included in the current array, and whether such corrections are applicable to other antigens necessitates empirical determination. Polyclonal corrections may be generalizable, however, the number of individual Abs that may simultaneously bind their cognate antigen is influenced by immunodominance of epitopes and steric hindrance, an idea supported by structural studies [29–31].

The high-dimensional assay we described greatly benefits from the use of the automated analytic platform. When combined, this system facilitates the rapid and accurate determination of antigen reactive Ab. Although the source material used as examples in this study are serum, the assay is amenable to many biological samples including nasal washes and saliva. The capacity to visualize both the magnitude and breadth of Ab responses to vaccination or infection with respiratory pathogens is valuable in determining correlates of protection derived from vaccination or pathogen exposure, and the ability to easily and rapidly expand the antigens incorporated into the array with minimal effects on the established analytics makes this approach an attractive option to fill the continuing need for large-scale serology studies.

## References

1. Garcia-Beltran, W.F., et al., Multiple SARS-CoV-2 variants escape neutralization by vaccine-induced humoral immunity. Cell, 2021. 184(9): p. 2372–2383 e9.

2. Yadav, P.D. and S. Kumar, Global emergence of SARS-CoV-2 variants: new foresight needed for improved vaccine efficacy. Lancet Infect Dis, 2022. 22(3): p. 298–299.

3. Aldridge, R.W., et al., SARS-CoV-2 antibodies and breakthrough infections in the Virus Watch cohort. Nat Commun, 2022. 13(1): p. 4869.

4. Alfego, D., et al., A population-based analysis of the longevity of SARS-CoV-2 antibody seropositivity in the United States. EClinicalMedicine, 2021. 36: p. 100902.

5. Hall, V.J., et al., SARS-CoV-2 infection rates of antibody-positive compared with antibody-negative health-care workers in England: a large, multicentre, prospective cohort study (SIREN). Lancet, 2021. 397(10283): p. 1459–1469.

6. Wang, Z., et al., mRNA vaccine-elicited antibodies to SARS-CoV-2 and circulating variants. Nature, 2021. 592(7855): p. 616-622.

7. Meyer, B., C. Drosten, and M.A. Muller, Serological assays for emerging coronaviruses: challenges and pitfalls. Virus Res, 2014. 194: p. 175–83.

8. Pisanic, N., et al., COVID-19 Serology at Population Scale: SARS-CoV-2-Specific Antibody Responses in Saliva. J Clin Microbiol, 2020. 59(1).

9. Caceres-Martell, Y., et al., Single-reaction multi-antigen serological test for comprehensive evaluation of SARS-CoV-2 patients by flow cytometry. Eur J Immunol, 2021. 51(11): p. 2633–2640.

10. Cameron, A., et al., A Multiplex Microsphere IgG Assay for SARS-CoV-2 Using ACE2-Mediated Inhibition as a Surrogate for Neutralization. J Clin Microbiol, 2021. 59(2).

11. Dawson, E.D., et al., Multiplexed, microscale, microarray-based serological assay for antibodies against all human-relevant coronaviruses. J Virol Methods, 2021. 291: p. 114111.

12. Egia-Mendikute, L., et al., Sensitive detection of SARS-CoV-2 seroconversion by flow cytometry reveals the presence of nucleoprotein-reactive antibodies in unexposed individuals. Commun Biol, 2021. 4(1): p. 486.

13. Guarino, C., et al., Development of a quantitative COVID-19 multiplex assay and its use for serological surveillance in a low SARS-CoV-2 incidence community. PLoS One, 2022. 17(1): p. e0262868.

14. Jolley, M.E., et al., Particle concentration fluorescence immunoassay (PCFIA): a new, rapid immunoassay technique with high sensitivity. J Immunol Methods, 1984. 67(1): p. 21–35.

15. Kenny, G., et al., Performance and validation of an adaptable multiplex assay for detection of serologic response to SARS-CoV-2 infection or vaccination. J Immunol Methods, 2022:16. p. 113345.

16. Roy, D.R., et al., Development, Validation, and Utilization of a Luminex-Based SARS-CoV-2 Multiplex Serology Assay. Microbiol Spectr, 2023: p. e0389822.

17. Stervbo, U., T.H. Westhoff, and N. Babel, beadplexr: reproducible and automated analysis of multiplex bead assays. PeerJ, 2018. 6: p. e5794.

18. Morgan, E., et al., Cytometric bead array: a multiplexed assay platform with applications in various areas of biology. Clin Immunol, 2004. 110(3): p. 252–66.

19. Allie, S.R., et al., The establishment of resident memory B cells in the lung requires local antigen encounter. Nat Immunol, 2019. 20(1): p. 97–108.

20. Beckett, D., E. Kovaleva, and P.J. Schatz, A minimal peptide substrate in biotin holoenzyme synthetase-catalyzed biotinylation. Protein Sci, 1999. 8(4): p. 921–9.

21. Nellore, A., et al., A transcriptionally distinct subset of influenza-specific effector memory B cells predicts long-lived antibody responses to vaccination in humans. Immunity, 2023.

22. fca, B.L. readfcs.m, Department of Medical Imaging, University of Debrecen. Version 22/Jun/2020. Matlab File Exchange. 2020; Available from: https://www.mathworks.com/matlabcentral/fileexchange/9608-fca_readfcs.

23. Finak, G., et al., Optimizing transformations for automated, high throughput analysis of flow cytometry data. Bmc Bioinformatics, 2010. 11.

24. G., C. L4P.m, Four parameters logistic regression - There and back again (2012). Matlab File Exchange. 2012; Available from: https://it.mathworks.com/matlabcentral/fileexchange/38122.

25. R., H. Flow Cytometry Data Reader and Visualization 2021; Available from: https://www.mathworks.com/matlabcentral/fileexchange/8430-flow-cytometry-data-reader-and-visualization.

26. Gaebler, C., et al., Evolution of antibody immunity to SARS-CoV-2. Nature, 2021. 591(7851): p. 639–644.

27. Henderson, R., et al., Controlling the SARS-CoV-2 spike glycoprotein conformation. Nat Struct Mol Biol, 2020. 27(10): p. 925–933.

28. Health, A.D.o.P. Coronavirus Disease 2019 (COVID-19). 2023; Available from: https://www.alabamapublichealth.gov/covid19/.

29. Barnes, C.O., et al., Structures of Human Antibodies Bound to SARS-CoV-2 Spike Reveal Common Epitopes and Recurrent Features of Antibodies. Cell, 2020. 182(4): p. 828–842 e16.

30. Nogal, B., et al., Mapping Polyclonal Antibody Responses in Non-human Primates Vaccinated with HIV Env Trimer Subunit Vaccines. Cell Rep, 2020. 30(11): p. 3755–3765 e7.

31. Han, J., et al., Polyclonal epitope mapping reveals temporal dynamics and diversity of human antibody responses to H5N1 vaccination. Cell Rep, 2021. 34(4): p. 108682.

