## Supplemental Figures for "A high-throughput multiplex array for antigen-specific serology with automated analysis"

**Figure S1.**

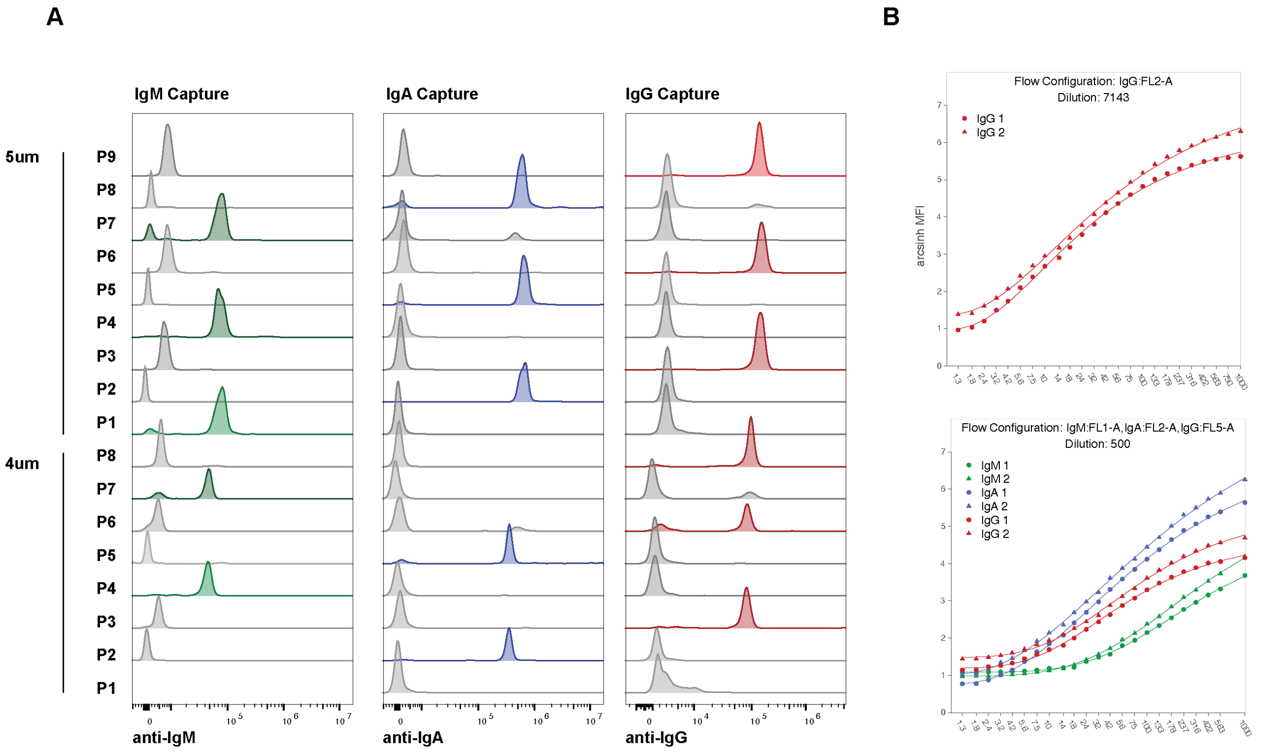


**Figure S1. Indirect standard curve generation and analysis**. A. The distribution of anti-Ig capture beads across the array constituents. All capture beads were mixed and stained with either IgG, IgM, or IgA and visualized with the indicated secondaries. **B.** Constructed standard curves from a batched run of samples. The software determines the number of dilutions used in a batch as well as the isotypes for the secondary antibodies for each dilution based on the specification in the sample manifest. The appropriate standard samples are automatically selected and 4PL curves are fit to data for each dilution, isotype and bead size. These fits are appropriately applied to convert transformed MFI into concentration units.

### Figure S2

A

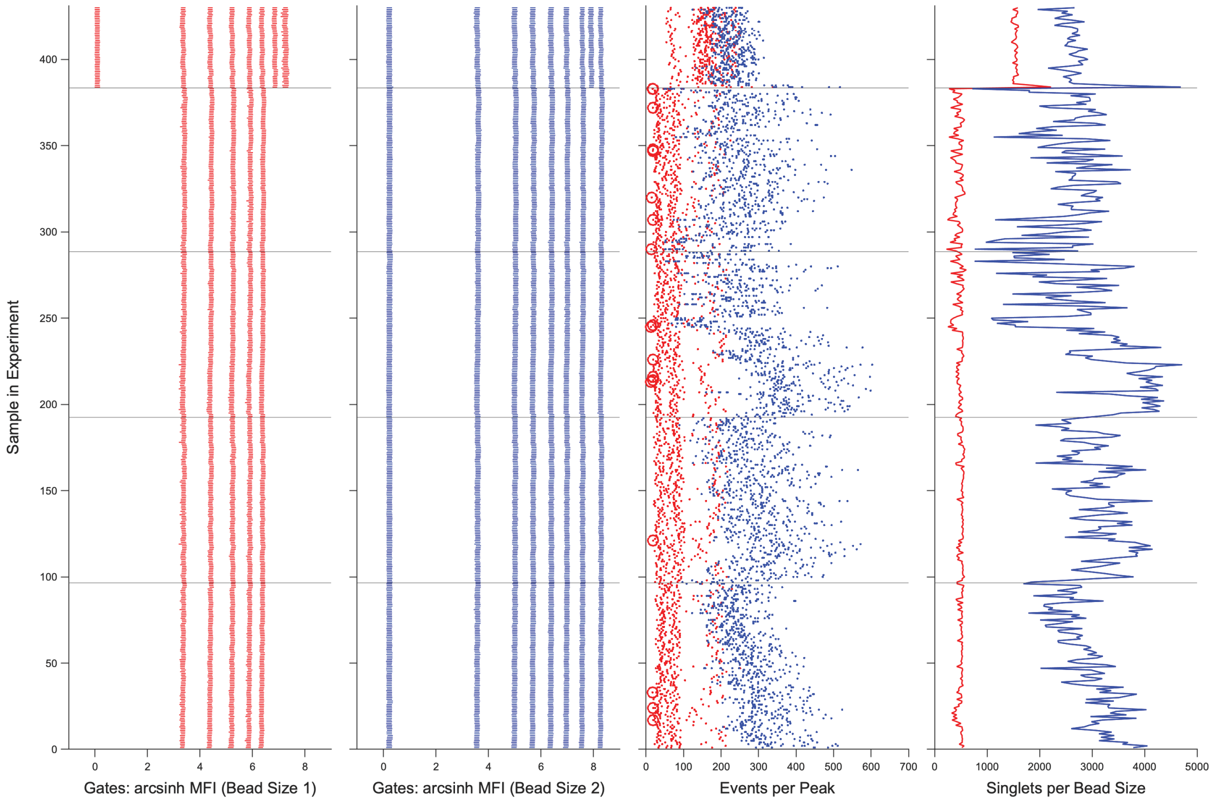

Sample in Experiment
Gates: arcsinh MFI (Bead Size 1)
Gates: arcsinh MFI (Bead Size 2)
Events per Peak
Singlets per Bead Size


B

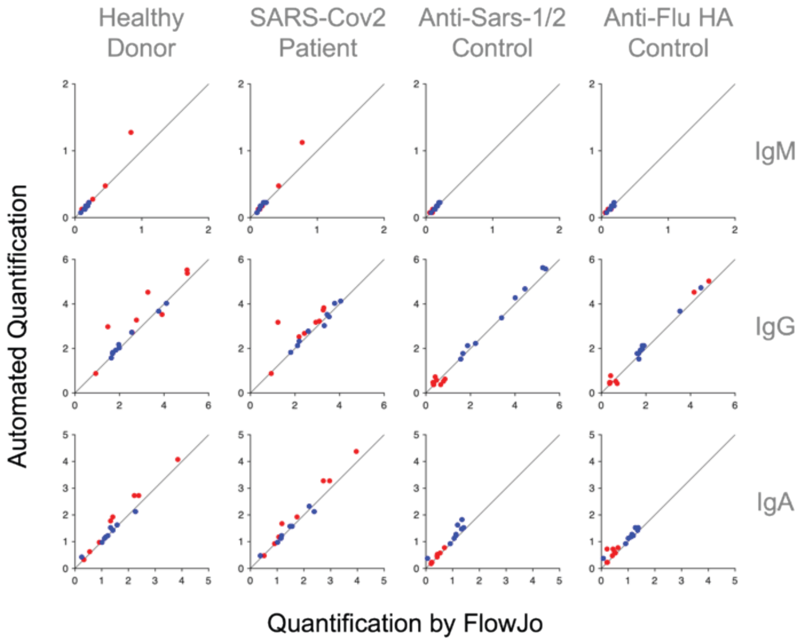

Healthy Donor
SARS-Cov2 Patient
Anti-Sars-1/2 Control
Anti-Flu HA Control
IgM
IgG
IgA
Automated Quantification
Quantification by FlowJo

**Figure S2. A. Quality assessment readout for batched run of samples.** For all plots, y-axis represents individual samples (FCS files) run from a manifest. Two leftmost plots show gate positions along the transformed MFI axis for 4um (left, red) and 5um (second from left, blue) beads. In this example, there were five 4um beads and eight 5 um bead gates for normal samples, and eight and nine 4um and 5um beads, respectively for the isotype standard samples. In this batch, there were two dilutions used per sample – 1:500 (for IgM, IgA and IgG) and 1:7000 (for IgG). Thus, there were 48 concentration standard samples – 24 for each dilution at the top of the plots. The third from left plot indicates the number of events per peak per sample (red, 4um beads; blue, 5um beads). Circles indicate beads that fall below a specified threshold (in this example, 20). The rightmost plot shows the number of singlets for 4um beads (red) and 5um beads (blue). **B.** Comparison of manual analysis (FlowJo) and automated analysis. Each column in the array of plots is a test sample and each row is an isotype. 4um and 5um beads are in red, blue, respectively. Automated analysis (y-axis) is in hyperbolic arcsine transformed MFI, and FlowJo analysis in in biexponential-transformed MFI.

**Figure S3**

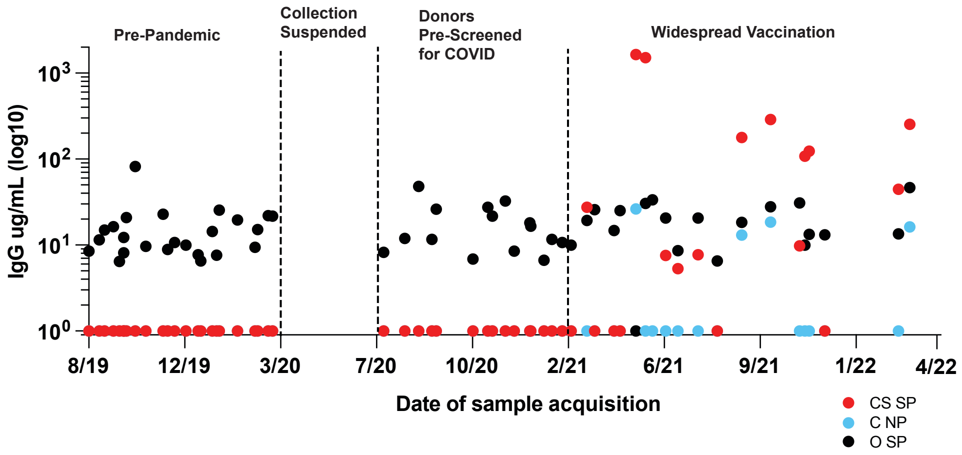


**Figure S3. The emergence of C SP reactivity during the SARS-CoV-2 pandemic.** The values of C SP, C NP, and O SP IgG for all donors plotted by collection date. The stages of the pandemic are highlighted.

#### Figure S4

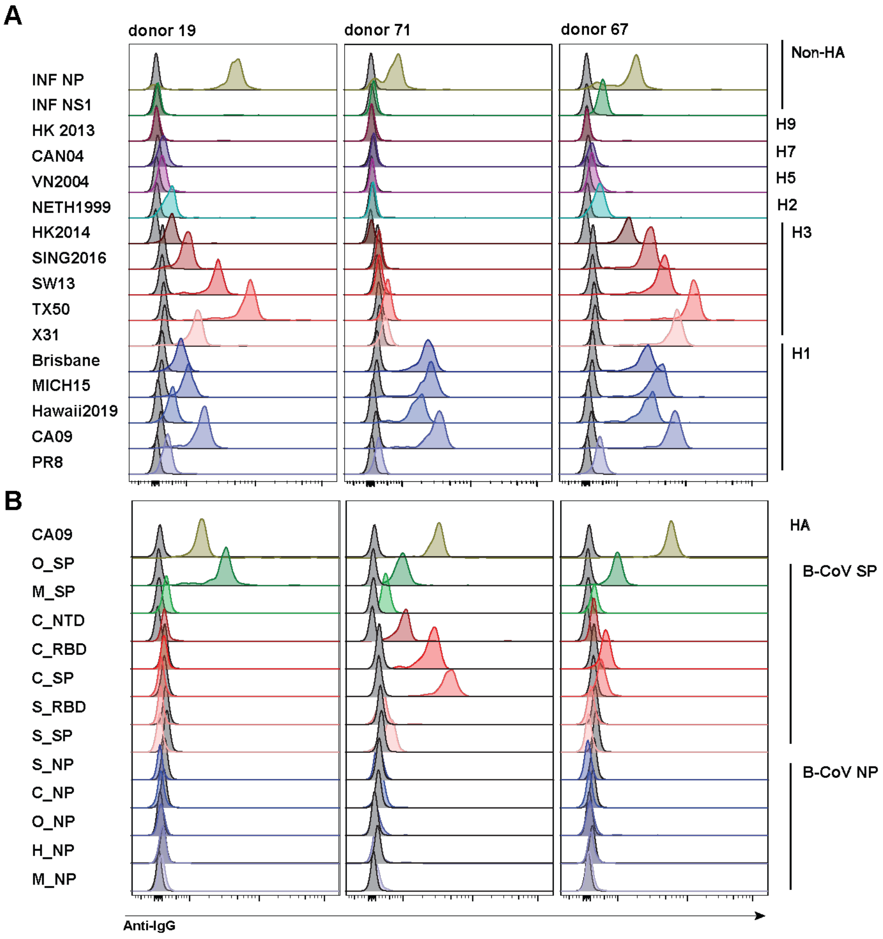


**Figure S4. Histograms of the antigen-conjugated beads stained with serum from the indicated donors and an anti-IgG secondary.** The Influenza array is shown in top panel, and the covid array is shown in the lower panes. Raw MFI derived from these flow files is used to calculate antigen specific IgG against each element of the array.
